# Multimodal Analysis of Sepsis-induced Cardiomyopathy in a Baboon Model

**DOI:** 10.1101/2025.06.16.659963

**Authors:** Tomohiro Abe, Robert Silasi, Ravi S. Keshari, Narcis Popescu, Girija Regmi, Satoshi Matsuzaki, Constantin Georgescu, Mariko Kudo, Cristina Lupu, Kenneth Humphries, Joe H. Simmons, Florea Lupu

## Abstract

Sepsis-induced cardiomyopathy (SIC) significantly contributes to sepsis-related morbidity and mortality, necessitating a deeper understanding of its mechanisms. This study used a post-hoc, multimodal approach—including single-nucleus RNA sequencing (snRNA-seq), echocardiography, mitochondrial function, and histopathology—to characterize SIC in a non-human primate model. Archived data and samples from six baboons challenged with 37.5 mg/kg of purified peptidoglycan were analyzed. Vital signs and echocardiography were monitored for 8 hours; the endpoint was survival at 168 hours or euthanasia for irreversible organ failure. Septic shock—defined by hypotension, tachycardia, and elevated lactate—was associated with poor outcomes. Echocardiography showed reduced intravascular volume, contraction, stroke volume, and cardiac index. SnRNA-seq revealed distinct transcriptomic profiles: non-survivors exhibited inflammation, mitochondrial dysfunction, and maladaptive remodeling; survivors showed activation of pathways supporting contraction, metabolism, and repair. Cell-type analysis highlighted metabolic dysfunction in cardiomyocytes, TNF/NF-κB-driven inflammation in endothelial cells, and stress responses in fibroblasts and pericytes. Mitochondrial analysis showed impaired electron transport and disrupted metabolism. Histopathology revealed inflammation and myofibrillar damage, more severe in non-survivors. This model recapitulates key SIC features and supports mechanistic and therapeutic discovery.

## INTRODUCTION

Sepsis is a critical condition caused by infection-induced dysregulation of the host responses, resulting in multiple organ failure and high mortality rates, especially in cases of septic shock ^1^. The primary pathophysiology of septic shock involves reduced intravascular volume and vascular resistance, resulting in decreased cardiac output and tissue perfusion. Despite advances in medical care, sepsis and septic shock still have a mortality rate of up to 50%, underscoring the urgent need for targeted treatments ^2, 3^.

Sepsis-induced cardiomyopathy (SIC) is recognized as a major contributor to morbidity and mortality ^4, 5^, accounting for up to 70% of cases ^6^. Although there are no definitive diagnostic criteria for SIC, common characteristics include systolic and diastolic dysfunction, ventricle dilations, and reversibility within 7–10 days ^7, 8^. Current guidelines suggest inotropic support for patients with persistent hypoperfusion and cardiac dysfunction despite adequate fluid resuscitation ^9^, however, the response to fluids and catecholamines may be reduced due to SIC ^7^. Several factors contribute to SIC progression, including oxidative stress, mitochondrial injury and dysfunction, impairment of intracellular calcium homeostasis, microcirculatory perturbation, and downregulation of beta-adrenergic receptors ^7, 8, 10^. Despite attempts to identify effective treatments for septic shock, none have consistently reduced mortality ^11^, highlighting the need for effective therapies.

Characterization of SIC in non-human primate (NHP) models is critical for understanding its pathology because rodent models do not accurately reflect human physiological responses ^12–14^. Peptidoglycan (PGN), a common component of the bacterial walls, has been shown to elicit potent inflammatory responses ^15^, similar to parental bacteria. We have successfully developed an NHP model of septic shock using PGN from *Bacillus anthracis* ^16, 17^. Given its highly inflammatory yet non-contagious nature, PGN emerges as a clinically relevant tool for SIC modelling.

We conducted a comprehensive post-hoc analysis on PGN-challenge experiments –including single-nucleus RNA sequencing (snRNA-seq) transcriptomic profiling, laboratory tests, cardiac functional assessments, mitochondrial functional assays and histopathological examinations– to characterize SIC in a NHP model of PGN-induced sepsis. Our findings provide a foundational framework to understand pathophysiology of SIC and exploring future therapeutic strategies to improve sepsis outcomes in humans.

Some of the results of these studies have been previously reported in the form of an abstract ^18^.

## RESULTS

### Survival and changes in vital signs after PGN-challenge

Three animals developed multiple organ failure and were humanely euthanized within six hours (non-survivors) while the other three survived until 7 days (survivors) (Fig. 1A). Mean arterial pressure decreased within 15 minutes and the hypotension was more profound and prolonged in non-survivors (Fig. 1B). Heart rate increased within 1 hour post-challenge, with no significant difference between the survivors and non-survivors (Fig. 1C).

**Fig. 1.**
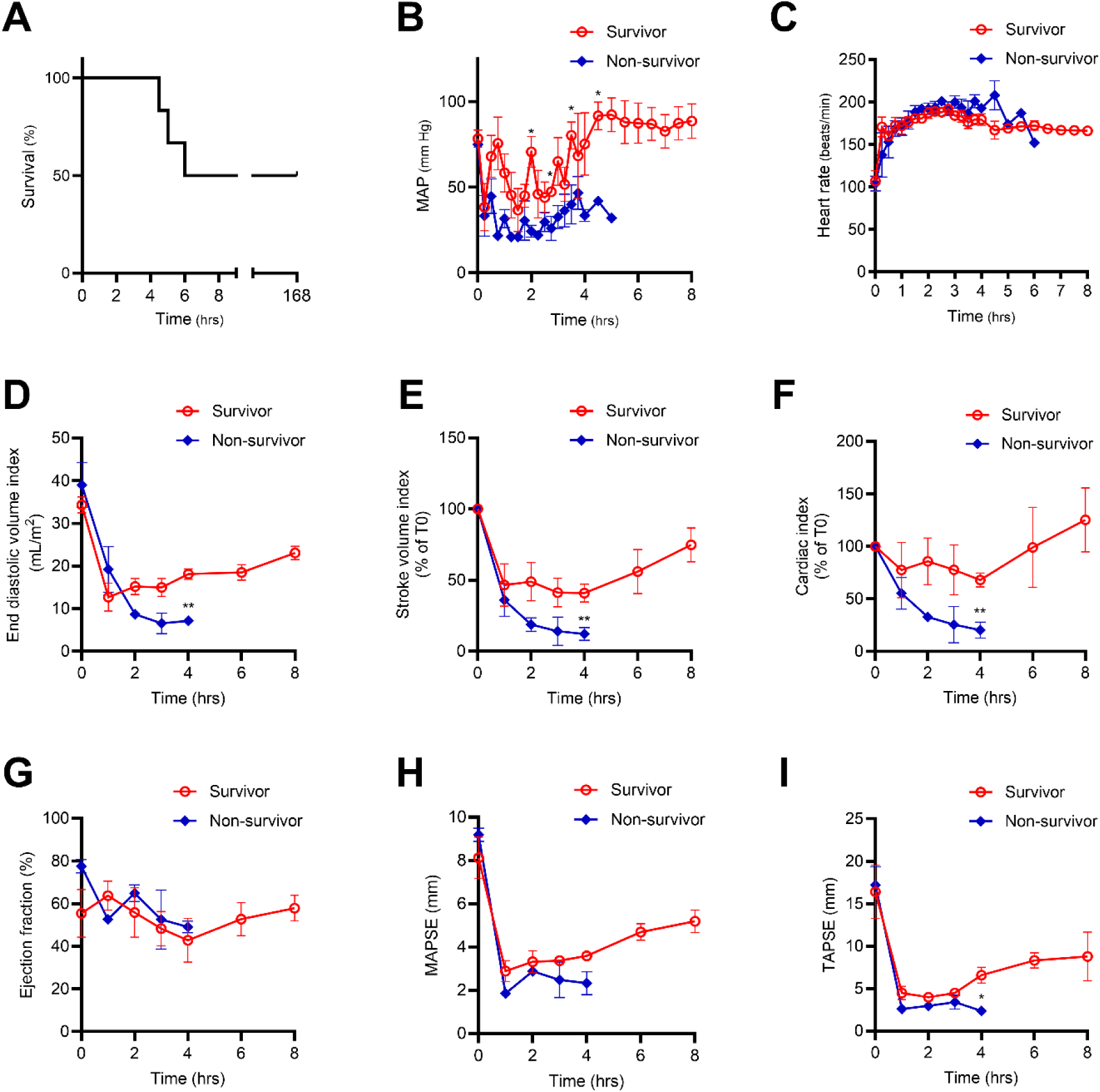
Survival rate and changes in vital signs and circulatory parameters following peptidoglycan challenge. (A) Survival rate after challenge. Changes in (B) mean arterial pressure (MAP), (C) heart rate, (D) the endo-diastolic volume index, (E) stroke volume index, (F) cardiac index, (G) ejection fraction, (H) mitral annular plane systolic excursion (MAPSE), and (I) tricuspid annular plane systolic excursion (TAPSE). For values where MAP (B) could not be measured due to severe hypotension, a value of 19 mmHg, below the lower limit of the device, was substituted. End-diastolic volume (D), stroke volume index (E), cardiac index (F) and ejection fraction (G) were calculated using echocardiographic measurements of the left ventricle diameters. Index values were normalized to estimated body surface area. Percent changes from baseline (T0) are shown for stroke volume index (E) and cardiac index (F). Values are shown as mean ±SEM. Group comparisons for each time point were performed using either a t-test or Mann-Whitney U test, depending on data distribution. *, p<0.05; **, p<0.01

### Echocardiography shows reduced intravascular volume and longitudinal contractility

The left ventricular end-diastolic volume index (EDVI) decreased abruptly (Fig. 1D), leading to corresponding decreases in stroke volume index (SVI) and cardiac index (CI). These reductions were more pronounced in non-survivors at T4 (Fig. 1, E and F). Short-axis systolic function, measured as ejection fraction (EF), did not significantly decrease (Fig. 1G). By contrast, the longitudinal-axis systolic function markers mitral annular plane systolic excursion (MAPSE) (Fig. 1H) and tricuspid annular plane systolic excursion (TAPSE) (Fig. 1I) dropped markedly at T1, and these changes were more severe in non-survivors; TAPSE reached statistical significance at T4 (Fig. 1I).

### SnRNA-seq reveals inflammation responses in non-survivors and recovery-related responses in survivors

We analyzed 13,425 nuclei from normal controls (4,586 nuclei), survivors (5,350 nuclei), and non-survivors (3,489 nuclei). In non-survivors, ∼1,500 genes were differentially expressed relative to controls, with marked upregulation of inflammatory (*GBP3, IL6, CXCL8*) and stress/injury genes (*HSP90AA1, HIF1A, CASP1*), and downregulation of extracellular matrix and integrin function (*COL10A1, COL3A1, PECAM1, LAMA2*), and cardiac muscle function genes (*ACTA1, STIM1, SCN7A, FGF14*) (Fig. 2A). In contrast, survivors exhibited fewer changes, with increased expression of genes supporting cardiac contraction (*ATP2A2, TPM1, FGF12, TNNT1, TNNI3*), mitochondrial function (*CHCHD10, ATP5F1A, SLC25A4, COX4I1*), and extracellular matrix organization (*DMD, COL1A1*), suggesting activation of recovery pathways (Fig. 2B).

**Fig. 2.**
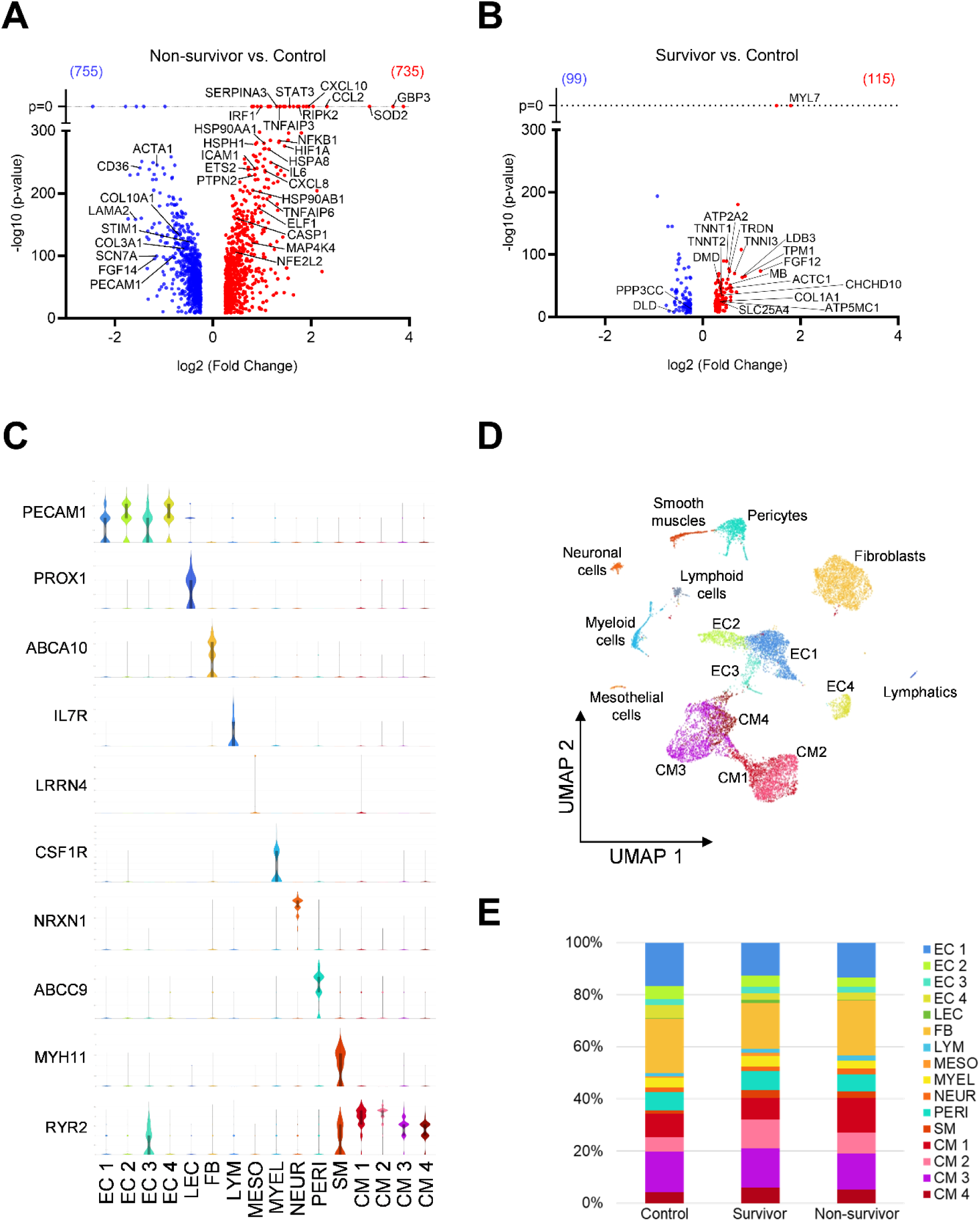
Cellular diversity, clustering, composition and differential expression in the myocardium revealed by snRNA-seq. (A, B) Volcano plots of differential expression genes (DEGs) of overall nuclei from (A) non-survivors and (B) survivors, compared to normal controls. DEGs with log2 (Fold Change)>0.25 and p<0.05 were plotted. (C) Violin plot of cell marker gene expression across clusters. (D) Uniform Manifold Approximation and Projection (UMAP) of myocardial 13425 nuclei, identifying 16 clusters, including four cardiomyocyte (CM1–4) clusters and four endothelial cells (EC1–4) clusters. (E) Comparison of cluster compositions across datasets.

Sixteen distinct cell clusters were identified based on specific marker genes (Fig. 2, C and D) including four cardiomyocyte (CM1–4) and four endothelial clusters—with similar distributions across groups (Fig. 2E).

*Cardiomyocyte* population consisted of 5,141 nuclei divided into four clusters: CM1 (1,327 nuclei), CM2 (1,126 nuclei), CM3 (2,003 nuclei), and CM4 (685 nuclei). Compared to controls, CM1–CM3 in non-survivors exhibited more differentially expressed genes (DEGs) than in survivors, while CM4 showed similar changes in both groups (Fig. 3A). IPA indicated activation of inflammatory signaling pathways and cellular stress/injury responses in all cardiomyocyte clusters of non-survivors (Fig. 3, B and C). In non-survivors, CM1, CM2 and CM3 showed downregulation of mitochondrial function (*COX6A2, COX6C, DLD, ATP5F1C*) and cardiac function genes. Differently, CM4 showed upregulation of several genes (*IDH2, ETFB, UQCRC1, CYC1, ATP5F1A*), key components of the mitochondrial energy-production machinery and redox regulation pathways. Upregulation of these CM4 genes is stronger in survivors suggesting this cluster contains cells undergoing an adaptive response aiming to sustain mitochondrial function, boost energy production, and mitigate metabolic stress in septic hearts.

**Fig. 3.**
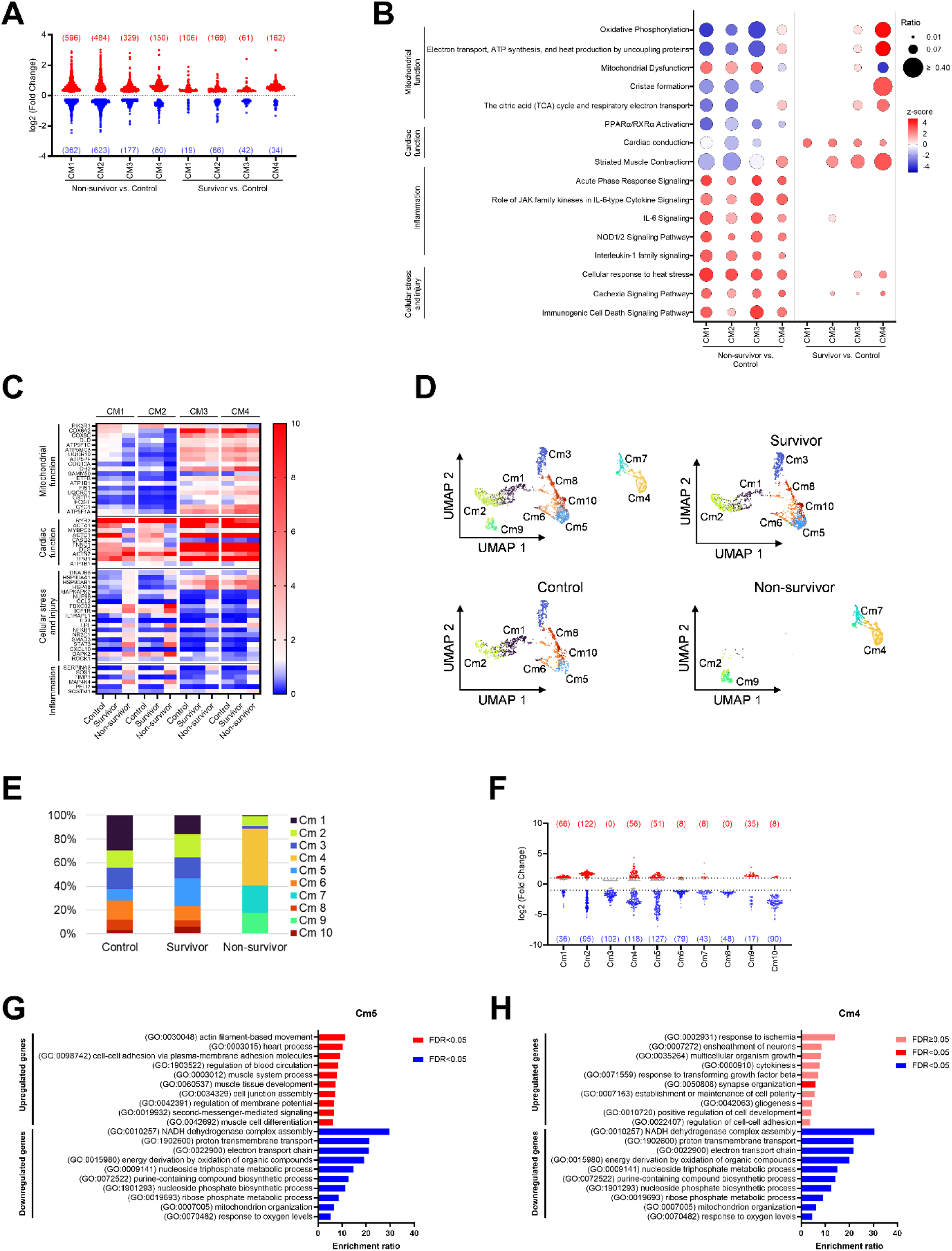
Transcriptomic analysis of cardiomyocyte clusters (CM1–4) in PGN-challenged baboon hearts. (A) Numbers of the differentially expressed genes (DEGs) in non-survivors and survivors compared to normal controls. Genes with p < 0.05 are shown. (B) Selected pathways related to mitochondrial function, cardiac function, inflammation and cellular stress and injury. (C) Heatmap of gene expression across cardiomyocyte clusters (CM1–4) and datasets, with colors representing the average expression per group within each dataset. (D) Uniform Manifold Approximation and Projection (UMAP) plots of unsupervised re-clustering of cardiomyocyte clusters. (E) Cluster composition across datasets. (F) Numbers of DEGs with p-value < 0.05. Red and blue indicate upregulated and downregulated genes, respectively|log2(fold change)| > 1. (G and H) Functional enrichment of DEGs in (G) Cm5 and (H) Cm4. The analysis was performed using a web-based enrichment tool ^42^. DEGs with -log10(p-value) > 5 and |log2(fold change)| > 1 was used for this analysis.

Unsupervised re-clustering of the whole cardiomyocyte dataset identified ten cardiomyocyte groups (Cm1–Cm10). As cells distribution, survivors resembled the controls, with slight increases in Cm5 and Cm10, whereas non-survivors show three unique clusters (Cm4, Cm7, Cm9) (Fig. 3, D and E). Among them, Cm5 and Cm4 show the highest numbers of DEGs (Fig. 3F and table S1). In Cm5 (common for survivors and control), genes central to mitochondrial metabolism (*UQCR10, ATP5F1B, COX8A*) were decreased, indicating a persistent suppression of mitochondrial functions in survivors. Conversely, sarcomeric genes (*LRRTM3, TENM2, MYH7B*) were upregulated, suggesting attempts to restore or remodel cardiac structure following sepsis-induced damage (Fig. 3G). Cm4, which is unique to non-survivors, also exhibited downregulation of oxidative phosphorylation, the TCA cycle, β-oxidation, and mitochondrial organization, indicating reduced metabolic capacity and weakened contractile machinery. Concurrently, genes associated with oxidative stress, ischemia, and fibrotic remodeling were upregulated, suggesting maladaptive responses (Fig. 3H).

*Endothelial cells* (EC) segregated into four major clusters. EC1 expressed genes specific to capillaries (*ROBO4, KDR, CD36*), EC2 to arterial ECs (*EFNB2, DLL4, HEY1/2, DKK2*), and EC3 to venous ECs (*NRP2, ESAM*). EC4 exhibited endocardial cell-specific gene expression (*NPR3, NRG1, BMP6*) (Fig. 4A). Capillary EC was the most abundant cluster, with similar proportions across datasets (Fig. 4B). In non-survivors, all four EC types were altered relative to controls, whereas in survivors most changes occurred in capillary ECs (Fig. 4C). In non-survivors, capillary EC exhibited upregulation of inflammatory response genes (*SOD2, GBP3, CXCL10, CCL2, HIF1A, IL6*) (Fig. 4D) and activation of TNF/NF-κB mediated inflammation pathways (Fig. 4, E and F). These changes led to enhanced functions in endothelial adhesion, mononuclear cell interaction, and antigen-presenting cell responses, along with suppression of PPARα/RXRα activation (Fig. 4G). The other EC types showed similar changes but with lower amplitude. Conversely, in survivors, only capillary EC were differentially changed, upregulating genes (*MYL7, CACNA2D1, MEF2C, SLC8A1, TNNC1*) related to cardiac hypertrophy signaling, oxidative phosphorylation and decreased mitochondrial dysfunction (Fig. 4, H and I). These changes were not associated with endothelial-to-mesenchymal transition, suggesting ECs may aid cardiomyocyte healing by producing myocardial-related proteins.

**Fig. 4.**
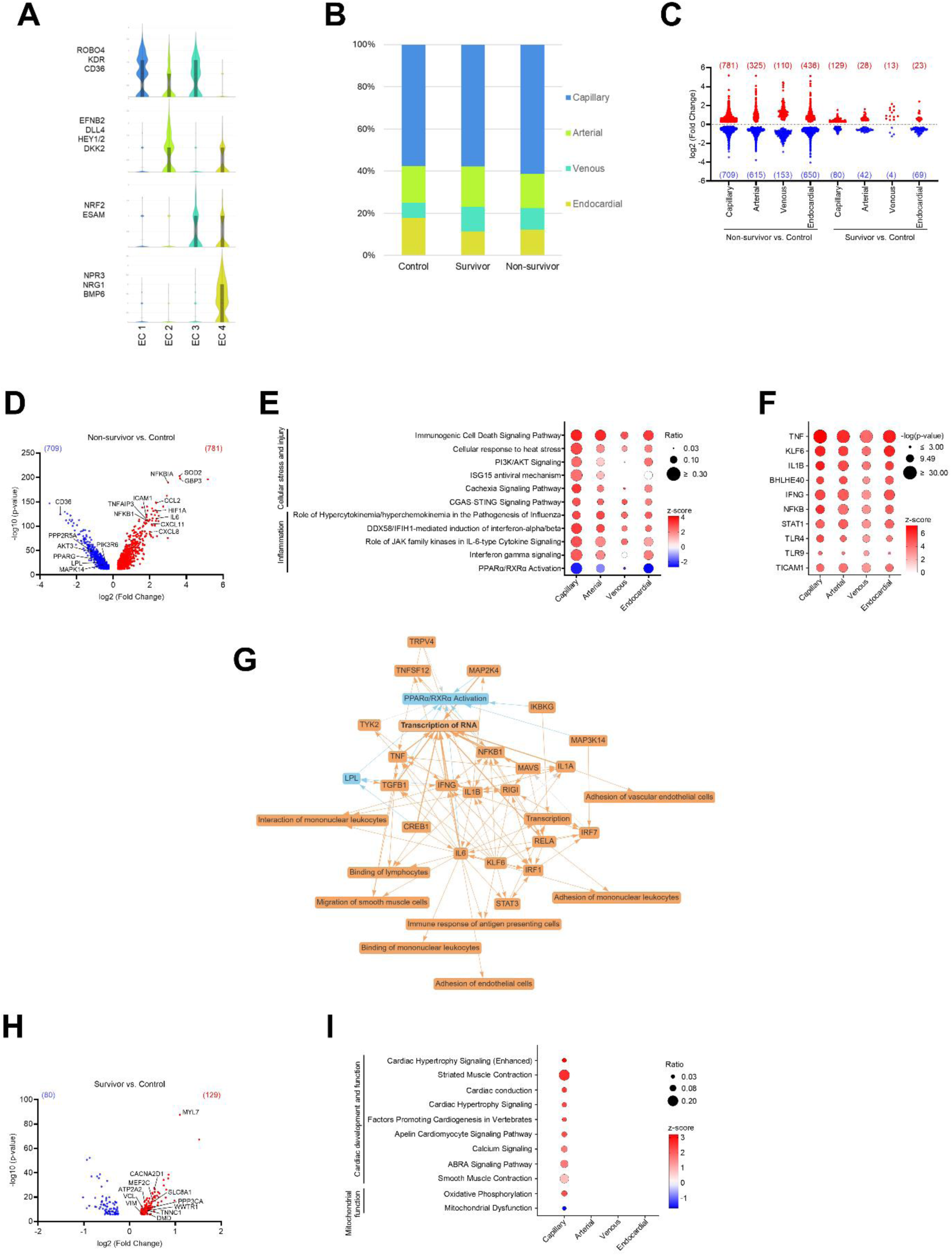
Transcriptomic analysis of endothelial cells (EC) (A) Violin plot showing expression of endothelial marker genes used to identify each EC cluster. (B) Composition of ECs across datasets. (C) Numbers of DEGs in non-survivors and survivors compared to normal controls. (D) Volcano plots showing DEGs of capillary ECs from non-survivors. |log2 (fold change)| > 0.25 and p-value < 0.05. (E) Canonical pathways and (F) upstream regulators identified in EC clusters from non-survivors. (G) Qiagen IPA graphical summary of capillary EC in non-survivors. Orange and blue indicate activation and inhibition, respectively; solid and dashed lines indicate direct and indirect effects, respectively. (H) Volcano plots showing DEGs of capillary ECs from survivors. (I) Canonical pathways associated with each EC cluster in survivors.

*Fibroblasts* (2,631 nuclei) and *pericytes* (994 nuclei) in non-survivors exhibited upregulation of inflammatory response genes (*GBP3*, *CCL2*, *STAT3*, *CXCL10*, *NFKB1*) (Fig. 5, A and B). IPA indicated activation of inflammatory pathways (interferon-gamma signaling, IL-6 signaling) and cellular stress responses (Fig. 5C). Genes in extracellular matrix organization pathways (*COL15A1, LAMA2, LAMC1, PBX1, ASPN, COL5A2*, *COL3A1*) were suppressed (Fig. 5, A to C). In survivors, fibroblasts showed limited gene expression changes, with some suppression of extracellular matrix organization due to downregulation of *LAMA2, LAMB1, LAMC1,* and *NTN4* (Fig. 5, A and C), while pericytes did not exhibit significant changes (Fig. 5, B and C). *Myeloid cells* (504 nuclei). In non-survivors, myeloid cells showed upregulation of immune activation genes, cytokines, chemokines, and hypoxia markers (*CD44, CXCL8, HIF1A*) (Fig. 5D) driving pro-inflammatory responses and immune cell adhesion (Fig. 5E). Survivors showed limited differential gene expression, with downregulation of genes related to myeloid cell functions (*DAB2, MS4A7, MERTK*) (Fig. 5D).

**Fig. 5.**
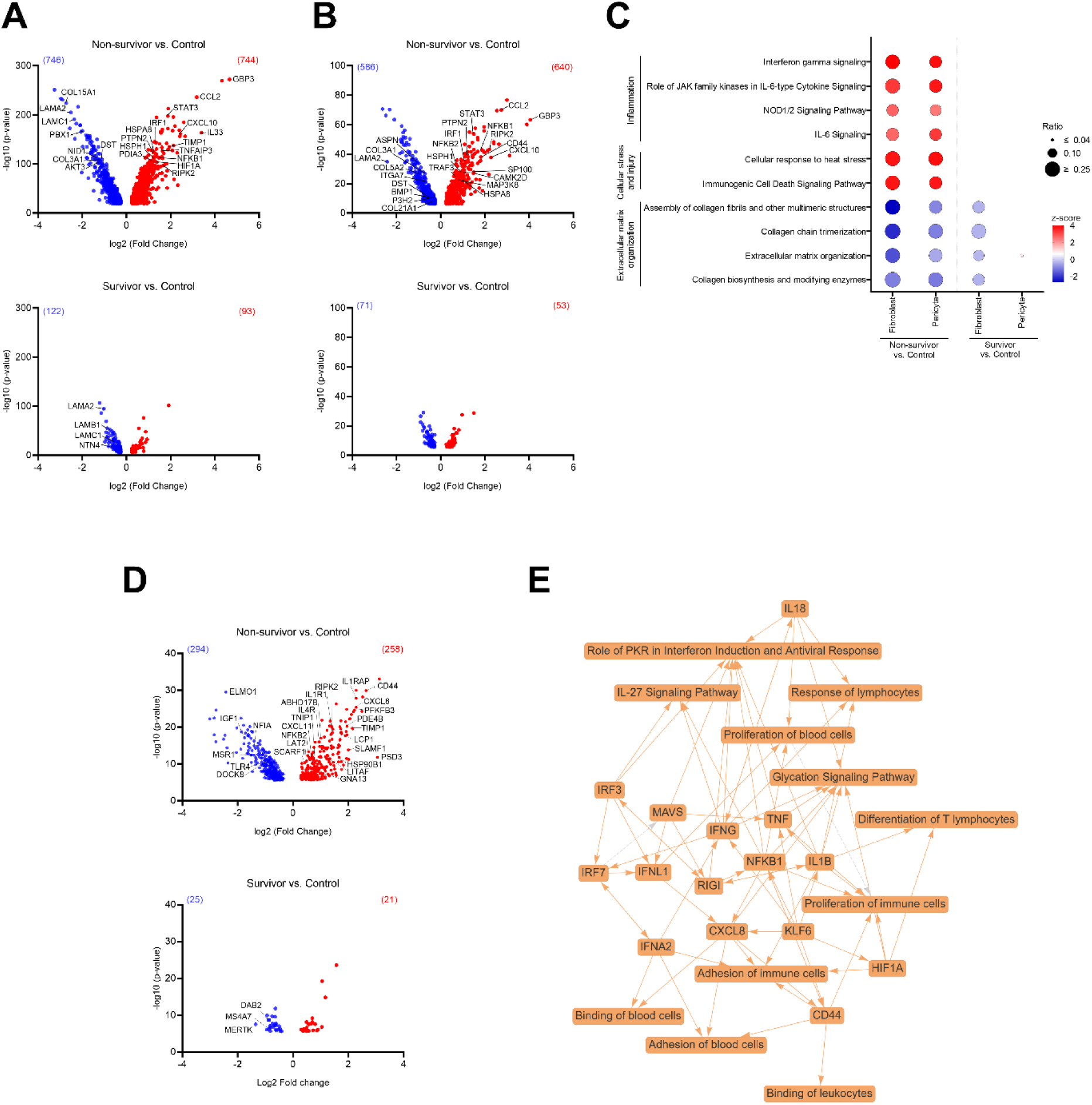
Transcriptomic changes in fibroblasts, pericytes and myeloid cells. Volcano plots showing DEGs in (A) fibroblasts and (B) pericytes from non-survivors and survivors compared to normal controls. Genes with |log2 (fold change)| > 0.25 and p-value < 0.05 are displayed. (C) Canonical pathways of fibroblasts and pericytes in non-survivors and survivors. The pathway names are displayed as listed in Qiagen IPA. (D) Volcano plots of DEG in myeloid cells from non-survivors and survivors compared to normal control. Genes with |log2 (fold change)| > 0.25 and p-value < 0.05 were plotted. (E) Qiagen IPA graphical summary of myeloid cells in non-survivors. Orange and blue indicate activation and inhibition, respectively. Solid and dashed lines indicate direct and indirect effects, respectively.

### Mitochondrial function is impaired in the heart muscle

PGN-challenge reduced nicotinamide adenine dinucleotide and hydrogen (NADH) oxidase, complex I, and complex III-IV activities in the heart (Fig. 6, A to C), indicating broad mitochondrial dysfunction that persisted throughout the experiment. While the pyruvate dehydrogenase (PDH) phosphorylation ratio (p-PDH/PDH) was not statistically significant, non-survivors consistently showed lower ratios, suggesting higher PDH activity (Fig. 6D). Carnitine palmitoyltransferase 1 (CPT-1) activity was also elevated in non-survivors compared to survivors and controls (Fig. 6E), indicating increased influx of oxidizable substrates into mitochondria.

**Fig. 6.**
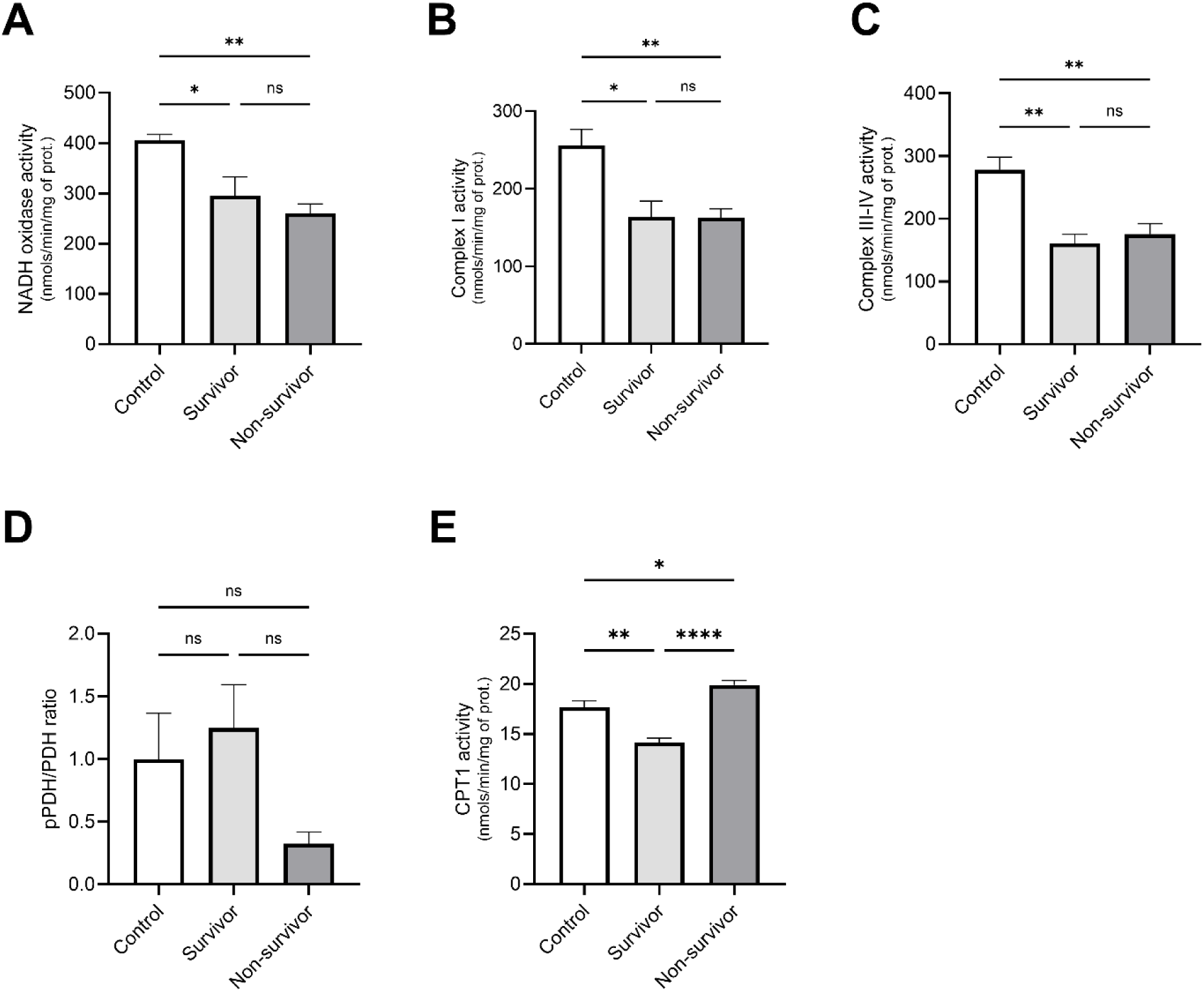
Mitochondrial function in PGN-challenged baboon hearts. (A) Nicotinamide adenine dinucleotide and hydrogen oxidase (NAD H) activity. (B) Complex I activity. (C) Complex III-IV activity. (D) Pyruvate dehydrogenase (PDH) ratio of phosphorylated to unphosphorylated (pPDH/PDH). (E) Carnitine palmitoyl transferase 1 (CPT 1) activity. Bars and whiskers represent mean ± SEM. Group comparisons were performed using Tukey’s or Dunn’s multiple comparisons test, depending on data distribution. *, p < 0.05; **, p < 0.01; ***, p < 0.001; ****, p < 0.0001; ns, not significant.

### Blood cells, coagulation and biochemical parameters

Hematocrit increased at T1 in both groups, with no significant difference between them (Fig. 7A). The lactate levels rose in both groups at T1, consistent with septic shock and the elevation was significantly higher in the non-survivors at T4 (Fig. 7B).

**Fig. 7.**
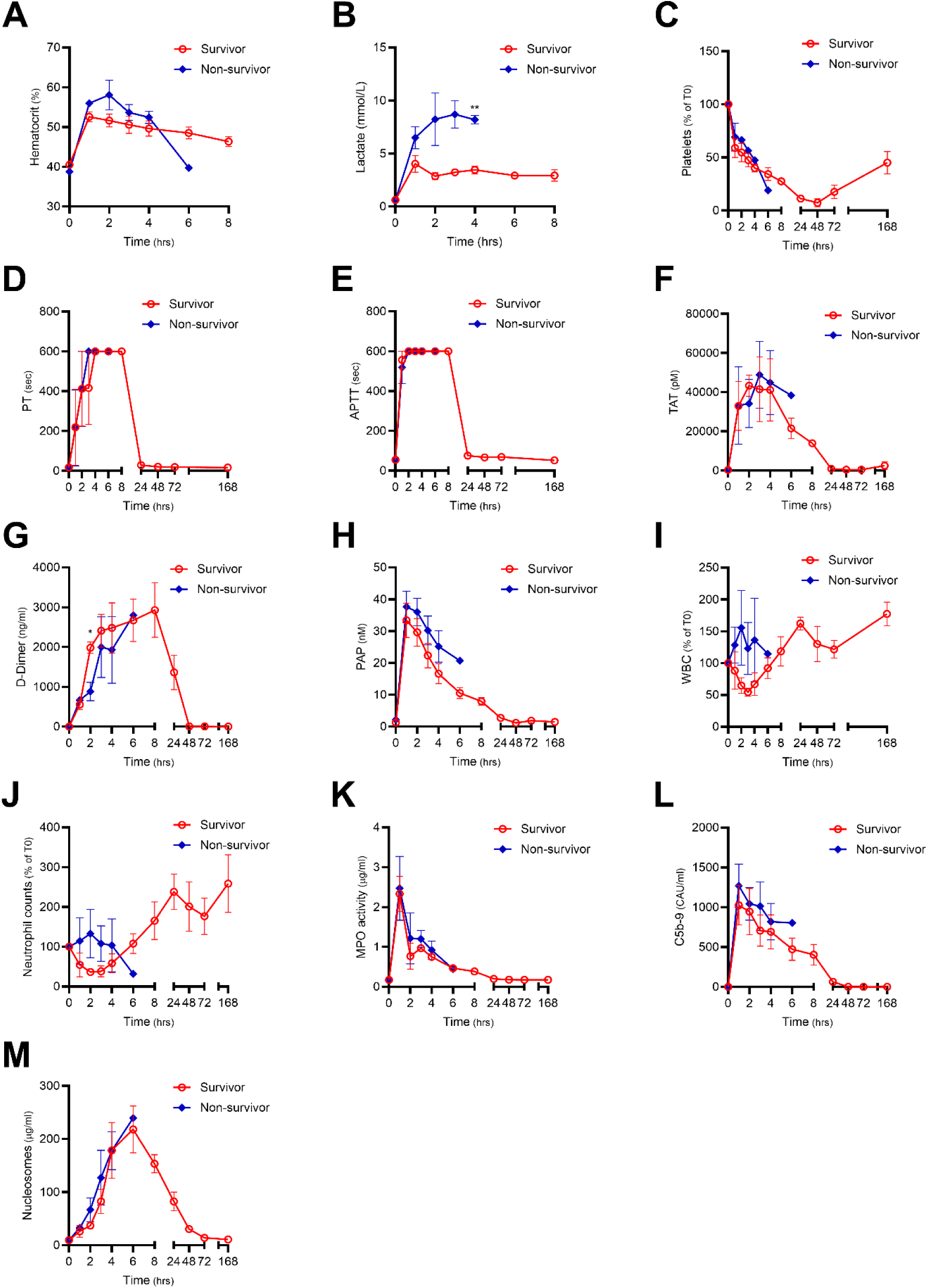
Changes in blood cells, coagulation parameters, and biochemical parameters after PGN-challenge. Shown are changes in (A) hematocrit, (B) lactate, (C) platelets counts, (D) prothrombin time (PT), (E) activated partial thromboplastin time (APTT), (F) thrombin-antithrombin complex (TAT), (G) D-dimer, (H) plasmin-α2-antiplasmin complex (PAP), (I) white blood cell counts, (J) neutrophil counts, (K) myeloperoxidase (MPO) activity, (L) sC5b-9, and (M) nucleosome. For platelet counts (C), white blood cell counts (I) and neutrophil counts (J), changes over T0 are shown. Values are presented as means ±SEM. Group comparisons for each time point were performed using t-test or Mann-Whitney U test, depending on data distribution. *, p < 0.05

Platelet counts decreased in both groups by 1 hour post-challenge and remained low throughout the observation period (Fig. 7C). Prothrombin time (PT) was prolonged after the challenge and exceeded the upper limit for all baboons at T4 (Fig. 7D). activated partial thromboplastin time (APTT) also became prolonged at T1 and surpassed the upper limit in all animals at T2 (Fig. 7E). thrombin-antithrombin (TAT) complex, D-dimer and plasmin-antiplasmin (PAP) complex levels all increased after PGN-challenge, with similar trajectories in both groups (Fig. 7, F to H).

White blood cells (WBC) showed divergent patterns - initially decreasing and then rising in the survivors and rising then decreasing in the non-survivors, though these trends were not statistically significant (Fig. 7I). Changes in neutrophil counts account for the changes in WBC (Fig. 7J). Myeloperoxidase (MPO) activity increased at T1 and subsequently decreased, without inter-group differences (Fig. 7K).

The soluble C5b-9 (sC5b-9) (an end-product of the complement activation), increased at T1 and then decreased in both groups, with no significant difference between them (Fig. 7L). Nucleosome levels increased gradually until T6, with no significant differences between the two groups (Fig. 7M). Overall, plasma coagulation changes, innate immune responses, and markers of systemic cell injury or death showed no statistically significant differences between survivors and non-survivors.

### Histopathological analysis evidenced myocardial inflammation and injury

HE-staining revealed interstitial edema in the myocardium of PGN-challenged baboons. Although milder in survivors, edema was observed in both groups (Fig. 8, A and B). Nuclear shrinkage with condensed chromatin, as well as microhemorrhages, were detected in non-survivors (Fig. 8A). Neutrophil infiltration was also evident in the myocardium of PGN-challenged baboon; although not extensive, it was still present in survivors (Fig. 8, A and C). These findings suggest that PGN challenge sustained inflammatory response and fibrin deposition in the myocardium over several days. Immunofluorescence showed fibrin/fibrinogen deposition in the myocardium of non-survivors but not in survivors (Fig. 8, A and D).

**Fig. 8.**
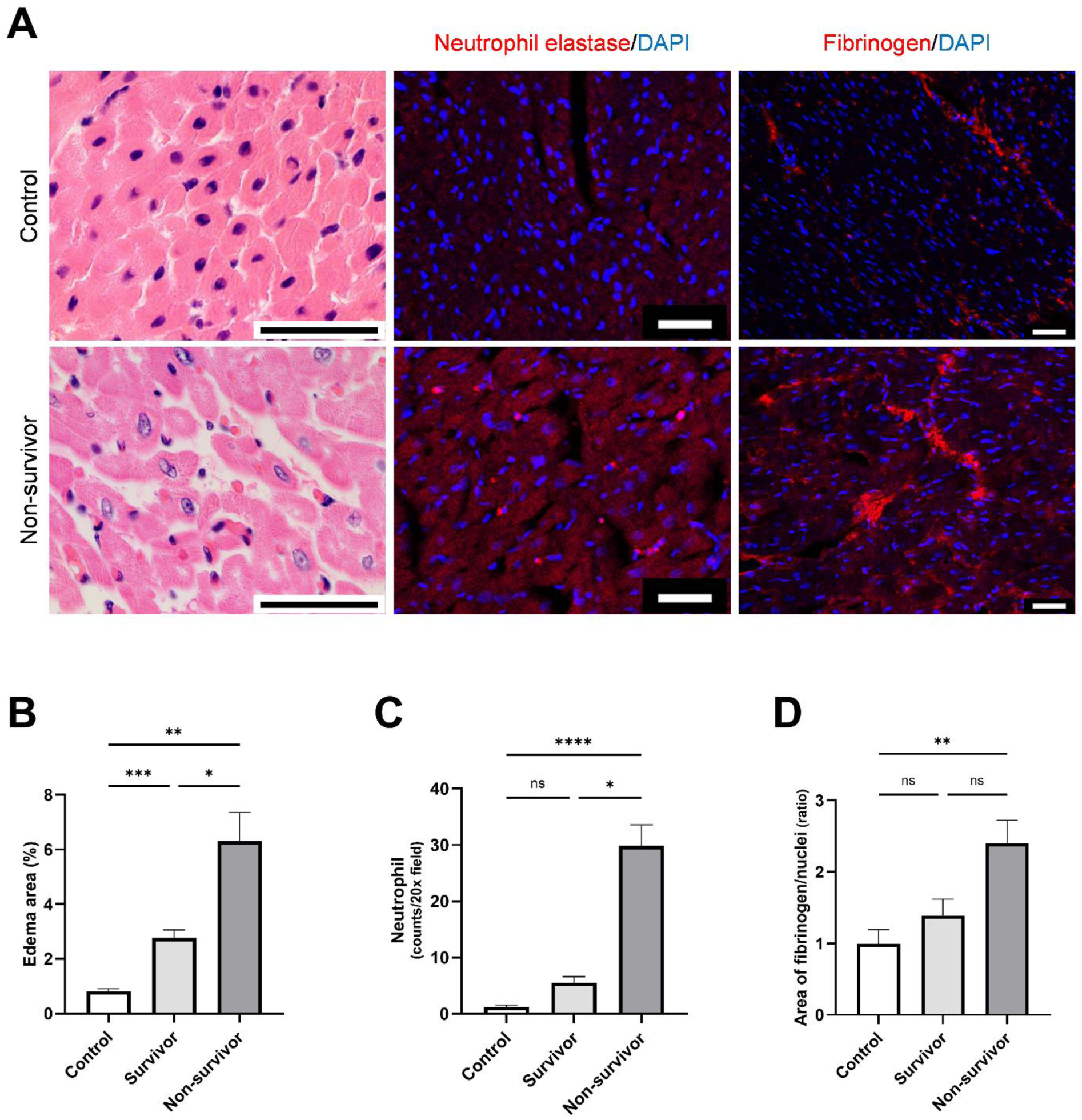
Histopathological analysis of myocardium in PGN-challenged baboon. (A) Hematoxylin and eosin (HE) staining and immunofluorescence staining for neutrophil elastase and fibrin/fibrinogen. Scale bars: 50 μm. Group comparison of (B) percentage edema (white) area in HE stained sections, (C) neutrophil count and (D) fibrinogen to nucleus ratio (n=10 microscopic fields). Bar and whiskers represent mean ±SEM. Group comparisons were performed using Tukey’s or Dunn’s multiple comparisons test, depending on data distribution. *, p < 0.05; **, p < 0.01; ***, p < 0.001; ****, p < 0.0001; ns, not significant.

## DISCUSSION

We analyzed cardiac-specific responses in baboons challenged with PGN—an abundant bacterial wall component mimicking Gram-positive sepsis ^16., 17^. PGN provoked severe hemodynamic instability, pronounced myocardial injury, and distinct transcriptomic shifts in multiple cardiac cell populations, all resembling human sepsis. Because non-survivors succumbed early, their data capture initial sepsis events, while survivors reflect more subacute changes of the heart. During early phase of fulminant sepsis, coagulation and innate immune pathways are vigorously activated, fueling widespread inflammation and cellular dysfunctions across multiple cell types in myocardium, on the other hand, these changes subside and recovery of cellular functions in the subacute phase.

This study provides a comprehensive analysis of sepsis-induced cardiac changes using single-nucleus RNA sequencing in a primate model, an approach not used yet in both primates and humans. Rodent models often fail to replicate key features of human septic cardiovascular failure ^12–14^, while human studies are limited by patient heterogeneity and scarce myocardial samples ^6–8^. Previously, myocardium from *S. pneumoniae*–challenged baboons revealed inflammatory changes and cell death linked to direct bacterial invasion ^19^. Our findings with sterile PGN infusion underscore that inflammation alone can drive sepsis-induced cardiomyopathy.

Our NHP model represents a fulminant form of sepsis, closely resembling the δ phenotype— characterized by septic shock, liver dysfunction, and high mortality ^20^. All animals experienced abrupt hypotension, more severe and prolonged in non-survivors, reflecting early septic shock seen in human patients. Concurrent decreases in EDVI, SVI, and CI further suggest reduced preload and impaired cardiac output. While classic SIC markers like reduced EF and ventricular dilation were absent, this likely reflects the study design (lack of fluid therapy) and sepsis heterogeneity. Contrary to the relatively stable short-axis EF, the longitudinal contractility, MAPSE and TAPSE, declined sharply, suggesting that reduced intravascular volume may mask early cardiac injury. The observed decline in longitudinal contractility aligns with evidence that longitudinal strain is highly sensitive to septic myocardial dysfunction ^21, 22^, underscoring the need to assess both short- and long-axis contractility in sepsis.

Laboratory indices confirmed severe sepsis. Platelet counts dropped rapidly, and prolonged PT and APTT reflected DIC-like changes. Lactate levels rose in both groups but were markedly higher in non-survivors, indicating severe tissue hypoperfusion and metabolic stress—hallmarks of poor outcomes. While innate immune markers (WBC, neutrophils, MPO, sC5b-9, circulating nucleosomes) were elevated in all animals, they did not predict survival, suggesting mortality was driven more by cardiovascular collapse and end-organ dysfunction than inflammation alone. Microscopic analysis revealed interstitial edema, nuclear shrinkage with chromatin condensation, microhemorrhages, and neutrophil infiltration, consistent with acute septic myocardial injury. Fibrin/fibrinogen deposition was prominent in non-survivors, highlighting the interplay of inflammation, coagulopathy, and hypoperfusion. Survivors showed milder edema and inflammation. Overall, PGN-induced sepsis provokes a severe inflammatory and coagulopathic myocardial insult, potentially fatal without adequate compensation.

SnRNA-seq identified 16 cell clusters—including cardiomyocytes, EC, fibroblasts, pericytes, and myeloid cells. Although cluster distributions were similar across normal, survivor, and non-survivors, gene expression diverged significantly. Non-survivors displayed pronounced inflammatory, stress, and injury-related pathways, alongside suppression of mitochondrial, extracellular matrix, and contractile genes—consistent with the notion that compromised bioenergetics that may underlie the marked decline in longitudinal function. contributes to organ failure in sepsis. By contrast, survivors displayed a more restrained transcriptomic response, involving mild upregulation of genes tied to cardiac and mitochondrial function, hinting at a partially adaptive or reparative process.

Among the four main cardiomyocyte clusters, those in non-survivors showed strong activation of inflammatory and stress-response pathways along with downregulation of mitochondrial and contractile genes, a combination that may underlie the marked decline in longitudinal function. Survivors’ cardiomyocytes, on the other hand, maintained or upregulated mitochondrial and contractile pathways that support myocardial function and recovery responses, potentially avoiding the worst metabolic derangements.

Our mitochondrial analyses showed decreased electron transport chain (ETC) enzyme activity in both non-survivors and survivors, indicating mitochondrial dysfunction during the acute phase and incomplete recovery in the subacute phase of sepsis. This suggests that decreased ETC activity alone does not predict survival. Non-survivors had lower p-PDH/PDH ratios and higher CPT-1 activity, indicating greater oxidizable substrate influx into mitochondria. In contrast, survivors showed higher p-PDH/PDH ratio and lower CPT-1 activity, suggesting reduced nutrient metabolism alongside decreased ETC activity—potentially regulating acetyl-CoA and acyl-CoA flux to limit reactive oxygen species production.

EC in non-survivors similarly exhibited marked upregulation of TNF, IL1β, and IFNγ signaling, consistent with microvascular dysfunction—a key driver of tissue hypoxia and organ failure in sepsis. Fibroblasts and pericytes reflected this pattern as well, with impaired matrix organization and heightened inflammatory responses. Myeloid cells showed strong upregulation of cytokines (*TNF, IL6*, and *IFNG*) some known to promote compensatory cardiac hypertrophy, pathological remodeling and eventual heart failure if unmitigated. Not only myeloid cells but also EC, fibroblasts, pericytes, and cardiomyocytes produce pro-inflammatory cytokines ^23–27^.

In survivors, these cell types had comparatively minimal transcriptional changes, and some subclusters displayed mild upregulation of genes related to cardiac development or function. Capillary EC appeared to bolster cardiomyocytes by expressing genes associated with cardiac and mitochondrial function. Such alterations may represent an “adaptive” endothelial response aiding cardiomyocyte survival.

Bulk transcriptomes from deceased human sepsis patients similarly show upregulation of *TNF, IL-1, NF-κB*, and hypoxia-related genes, with concomitant downregulation of mitochondrial and M-band signaling ^9, 28, 29^. Rodent data further implicate inflammation, mitochondrial dysfunction, and pyroptosis in SIC ^29–31^. Our snRNA-seq data extend these insights, revealing that severe inflammation in cardiomyocytes compromises mitochondrial integrity, while endothelial and stromal cells also undergo stress and lose extracellular matrix support early in sepsis.

Our echocardiographic assessments revealed notably reduced intravascular volume, stroke volume, and cardiac index in non-survivors, underscoring the importance of maintaining adequate volume status during septic shock. Current sepsis guidelines recommend fluid resuscitation, and for patients who remain hypotensive despite sufficient fluids, inotropic support is often considered ^9^. However, inotropes carry heightened mortality risks ^32^, potentially through downregulation of β1-adrenergic receptor ^33^ and exacerbation of inflammation markers ^34^. Consequently, mechanical circulatory support has been proposed as an alternative approach for severe SIC ^29, 35^. Future therapies might also exploit anti-inflammatory strategies, including sphingosine-1-phosphate receptor agonists ^22^, and mitochondrial-targeted antioxidants ^36, 37^.

Overall, our findings highlight the multifaceted nature of sepsis-induced cardiomyopathy, encompassing systemic inflammation, coagulopathy, mitochondrial injury, and cellular stress. They also reaffirm the importance of nuanced cardiac function metrics (MAPSE, TAPSE) in sepsis. Utilizing an NHP model provides a valuable preclinical platform for targeting therapies aimed at preserving mitochondrial integrity, dampening maladaptive inflammation, and stabilizing cell functions.

Several limitations warrant consideration. First, we used data and tissues from experiments not originally designed for this purpose, and the protocol lacked standard shock treatments (e.g., fluid resuscitation or vasopressors). While this constrains direct extrapolation to clinical sepsis, it repurposed existing resources and avoided additional animal use. Second, single-nucleus RNA sequencing only captures nuclear transcripts, potentially missing certain isoforms or splicing events, but it enables analysis of bio-banked tissues and is less prone to artifacts than single-cell dissociation. Third, the PGN-challenge model may not fully reflect the complexity of sepsis, which often involves polymicrobial infections and comorbidities. Lastly, although NHPs mirror human inflammatory processes better than rodents, species-specific differences remain. Future validation of DEGs and pathways in human sepsis samples (e.g., patient cardiac biopsies or blood-based biomarkers) will confirm these findings and enhance their translational relevance.

In conclusion, PGN-induced sepsis in baboons recapitulates key features of septic shock and SIC, including acute hypotension, reduced intravascular volume, and impaired longitudinal cardiac function. Our molecular, transcriptomic, and mitochondrial analyses highlight heightened inflammation, mitochondrial dysfunction, and cellular stress as drivers of severe myocardial injury. Although this model does not fully capture the pathophysiology of clinical SIC, it offers a robust framework for investigating fundamental disease mechanisms and developing new therapeutic strategies to reduce sepsis mortality.

## MATERIALS AND METHODS

### Animal experimentation

We analyzed data and samples from historical controls of PGN-challenge experiments that were approved by the Institutional Animal Care and Use Committee at MD Anderson Cancer Center. All experiments were conducted in compliance with the Animal Welfare Act, the Guide for the Care and Use of Laboratory Animals, and the National Institutes of Health Office of Laboratory Animal Welfare. The experimental design and purification of PGN have been described previously ^16, 17^. Six *Papio anubis* baboons (7.4–10.3 kg) were anesthetized and fitted with venous catheters for administration and blood sampling. Each received a 15-minute intravenous infusion of purified PGN (37.5 mg/kg in saline), with T0 marking infusion start and survival assessed at 168 hours. Vital signs were recorded every 15 minutes for 8 hours before the animals were returned to their cages and monitored until 168 hours. Blood sampling and echocardiography were performed at T0, 1, 2, 3, 4, 6, 8, 24, 48, 72, and 168 hours. Animals developing irreversible multiple organ failure or reaching the 7-day end-point were humanely euthanized, according to the American Veterinary Medicine Association Panel on Euthanasia^38^. Tissues were collected for histopathological, transcriptomic, and mitochondrial analyses. Control biobanked tissues from healthy baboons (n=3) were used for comparison.

### Echocardiography

Echocardiographic measurements were performed using the handheld Butterfly iQ ultrasound (Butterfly Network, Inc., NY, USA). Left ventricular (LV) systolic and diastolic diameters were measured in the parasternal long-axis view, while mitral annular plane excursion (MAPSE) and tricuspid annular plane excursion (TAPSE) were assessed in the apical 4-chamber view ^39^. LV end-diastolic volume, ejection fraction, stroke volume, and cardiac output were calculated from LV diameters ^40^ and normalized to body surface area estimated from body weight ^41^.

### Single-nucleus RNA sequencing (snRNA-seq)

*Sample preparation and nuclei isolation*. Heart tissues stored at -80°C were used for snRNA-seq. Frozen heart tissue was cut into small pieces on a glass petri dish placed on dry ice, ensuring the sample remained frozen, and then quickly transferred to a pre-chilled C tube containing lysis buffer. The lysis buffer was prepared using Miltenyi Biotec nuclei extraction buffer supplemented with an RNase inhibitor (final concentration: 0.2 U/µL). Tissue dissociation was performed using the gentleMACS Octo Dissociator with heaters (Miltenyi Biotec) by selecting the preloaded gentleMACS Program 4C_nuclei_1. The resulting nuclei suspension was filtered through a 70 µm MACS SmartStrainer (Miltenyi Biotec) placed on a chilled 15 mL tube to remove tissue debris. To maximize nuclei recovery, the SmartStrainer was rinsed with ice-cold lysis buffer. The filtered suspension was centrifuged at 300×g at 4°C for 5 minutes. The supernatant was carefully aspirated, and the nuclei pellet was resuspended in ice-cold resuspension buffer by gently pipetting up and down 10 times using wide-bore pipette tips. To ensure removal of aggregates and debris, the nuclei suspension was filtered twice through 30 µm MACS SmartStrainers placed on chilled 15 mL tubes. Nuclei were then stained with AO/PI dye and counted using the K2 Cellometer (Nexcelom Bioscience). A total of 16,500 nuclei per sample were loaded to achieve an estimated target capture of 10,000 nuclei.

*Library preparation and sequencing.* Libraries were generated using the Chromium Next GEM Single Cell 3’ Reagent Kits v3.1 (Dual Index) (10x Genomics, cat#1000268), following the manufacturer’s protocol. Single-nucleus suspensions were prepared according to the sample manifest and encapsulated using the Chromium Controller (10x Genomics) to generate Gel Beads-in-Emulsion (GEMs), enabling barcoding of individual nuclei. Post-GEM generation, reverse transcription (GEM-RT) was performed within each GEM, followed by cleanup and cDNA amplification. The amplified cDNA was then fragmented, end-repaired, and A-tailed. Dual index adapters were ligated to the cDNA fragments, followed by PCR amplification. Library quantification was performed using a Qubit Fluorometer (ThermoFisher), and the size distribution was assessed using an Agilent 2100 Bioanalyzer (Agilent). Sequencing libraries were prepared to achieve a target of 350 million paired-end reads (700 million total), ensuring a coverage of 35,000 reads per nucleus for a targeted capture of 10,000 nuclei. Sequencing was conducted on an NovaSeq 6000 (Illumina) following the manufacturer’s recommendations.

*Quality control, filtering, and clustering.* The Cell Ranger (ver. 7.1.0, 10x Genomics) mkfastq analysis pipeline was used to demultiplex raw base call (BCL) files generated by the Illumina sequencer into FASTQ files. The cellranger count pipeline was then used to process these FASTQ files, performing alignment, filtering, barcode counting, and UMI counting. Feature-barcode matrices were generated using the Chromium cellular barcodes to determine clusters and conduct gene expression analysis. The filtered feature-barcode matrix from Cell Ranger was used for downstream analysis. Read count matrices were analyzed using the Seurat package in R. The feature-barcode matrix was imported into R using the Read10X function and converted into a Seurat object, followed by quality control (QC) filtering and normalization. For quality control, nuclei were filtered based on the following criteria: (a) doublets removal; (b) number of unique genes per nucleus: 500 < nFeature_RNA < 5000; (c) total number of molecules per nucleus: 1000 < nCount_RNA < 20,000; (d) mitochondrial genome reads: <30%. High-quality nuclei from all baboon types were integrated into a single dataset using the SCTransform data integration pipeline in Seurat with default settings. Highly variable features were selected, and dimensionality reduction was performed using principal component analysis (PCA) followed by uniform manifold approximation and projection (UMAP) and stochastic neighbor embedding (sSNE) via the RunUMAP function, using the top 3000 most variable genes. Nucleus clustering was conducted using the FindClusters function with a resolution of 0.015, applying the K-nearest neighbor (KNN) approach and smart local moving (SLM) modularity optimization. The resulting clusters were visualized in UMAP 2D representations. Cell type annotation was performed using the *celldex* R package for mixed cell deconvolution, referencing the adult Heart_HCA expression profile matrix obtained via the download_profile_matrix function in the SpatialDecon package. The dominant nuclear type within each cluster was assigned as the cluster cell type identity. Identification of conserved cell type markers within each cluster was performed using the *FindConservedMarkers* function in Seurat, while differential expression analysis between baboon phenotype groups was conducted using the *FindMarkers* function with default parameters. Genes were considered significantly differentially expressed (DEGs) if they met the following criteria: (a) adjusted p-value < 0.05; (b) absolute log2 fold change (|log2FC|) > 0.25. For visualization, Loupe Browser (ver. 8.0.0) was used to generate UMAP plots, violin plots, and conduct unsupervised re-clustering. Ingenuity Pathway Analysis (IPA) software (Qiagen) was used for biological pathway and functional analysis. Web based enrichment tool ^42^ was used for functional enrichment analysis of re-clustered cardiomyocyte clusters. The snRNA-seq datasets have been deposited in the Gene Expression Omnibus (GEO), accession number GSE292370.

### Mitochondrial functional analysis

Mitochondrial functions were assessed by electron transport chain (ETC) enzyme activities, carnitine palmitoyltransferase 1 (CPT-1) activity, and phosphorylation status of pyruvate dehydrogenase (PDH). As previously described ^43^, mitoplasts were isolated from frozen heart tissues (∼100 mg). Thawed heart tissues were minced in 5 mL of mitochondria isolation buffer containing 210 mM mannitol, 70 mM sucrose, 1.0 mM EDTA, and 5.0 mM MOPS pH 7.4, and homogenized by a motor-driven Potter-Elvehjem tissue grinder. The homogenate was spun at 750g for 5 min at 4°C, and the supernatant was then spun again at 10,000g for 15 min. The resulting mitoplasts pellet was resuspended in 25 mM MOPS pH 7.4 and immediately snap-frozen.

NADH oxidase, an overall assessment of electron transport chain activity through complexes I– III–IV was measured as follows. Twice frozen-thawed mitoplasts were diluted into 25 mM MOPS pH 7.4 at 25 µg/mL and the rotenone-sensitive rate of NADH oxidation was monitored spectrophotometrically utilizing an Agilent 8453 diode array UV−Vis spectrophotometer. Activity was measured as the rate of NADH oxidation (340 nm, ε = 6200 M^−1^ cm^−1^) following addition of 10 mM KCl and 150 μM NADH. Complex I activity was measured spectrophotometrically as the rate of NADH oxidation in the presence of 50 nM antimycin A and 100 μM ubiquinone-1 following the addition of 150 μM NADH. Complex III-IV combination activity was measured spectrophotometrically as the rate of ubiquinol oxidation (280 nm, ε = 13700 M^−1^ cm^−1^) after adding of 10 mM KCl and 100 μM ubiquinol-1 ^43^. The specificities for complex I and complex III-IV activities were confirmed by inhibition with 1.0 µM rotenone and 0.5 µM antimycin A, respectively. CPT-1 activity was assessed as the rate of carnitine-dependent CoA-SH formation from palmitoyl-CoA as previously described ^44^. The assay was performed in 25 mM MOPS pH 7.4 buffer containing 1 mM EGTA, and BSA (0.1%). Mitoplasts samples at 50 µg/mL were mixed with 100 µM palmitoyl-CoA and 100 µM DTNB, and incubated for 20 min at RT to purge residual endogenous carnitine in the samples. CPT1 activity was measured spectrophotometrically as the rate of increase at 412 nm upon the addition of carnitine (10 mM final). Each set was run in triplicate and with the inclusion of one sample assayed without carnitine, which was subsequently used to subtract the background, carnitine-independent, activity. We assessed phosphorylation status of PDH complexes (pho-PDH/PDH) as an indicative of endogenous activities by utilizing Western blot ^44^. Mitoplasts samples at 50 µg/mL were diluted with 4x NuPAGE sample buffer (Invitrogen) containing 25 mM DTT and heated at 95°C for 5 min. Western blot was performed following Li-COR manufacturer’s protocol. SDS-PAGE (4–12% NuPAGE BisTris gel, Thermo Fisher Scientific) gels were transferred to nitrocellulose membranes and blocked for 60 min with Odyssey TBS blocking buffer (LI-COR) at RT. Intensities of bands were standardized to the total protein loaded (Revert Total Protein Stain; LI-COR, 926–11014). Antibodies used were Cell Signaling product nos. 3205S and 31866S for PDH and pho-PDH (Ser293), respectively. Antibodies were diluted 1:1000 in block buffer added overnight at 4°C and subsequently washed the following day, and the secondary antibody (IRDye 800CW, LI-COR; 1:2,000 dilution) was incubated for 1 h. Following additional washing, blots were analyzed on an Odyssey CLx imaging system using Image Studio software (LI-COR). The protein concentration was determined by the BCA method (Thermo Scientific) with a BSA as the standard.

### Laboratory tests

Complete blood counts were measured with a VetScan HM5 analyzer (Abaxis Inc, CA, USA). Lactate was measured by Lactate Plus Meter (Nova Biomedical Corp. MA, USA).

### Measuring plasma markers of coagulation, innate immune responses and cell death

PT and APTT were measured as described ^45^. Plasma concentrations of TAT complex, D-dimer, and PAP complex, sC5b-9, MPO activity and nucleosome were measured as described ^46^.

### Histopathological analysis

Formalin-fixed, paraffin-embedded heart tissues were stained with hematoxylin and eosin (HE). Cryo-embedded tissue sections were stained for neutrophil elastase and fibrin/fibrinogen, with nuclei counterstaining ^46^. Images were analyzed using ImageJ (version 1.54f, NIH, Bethesda, MD, USA).

### Statistical analysis

All statistical analyses, except for snRNA-seq data, were performed using Prism (ver. 10.2.3, GraphPad Software, Boston, MA). The normality of continuous variables was assessed with the Shapiro–Wilk test. Depending on the number of groups, the data distribution, and whether data were paired or unpaired, Mann–Whitney U test, Wilcoxon matched-pairs signed-rank test, one-way analysis of variance (ANOVA) with Tukey’s multiple comparison test, or Kruskal–Wallis test with Dunn’s multiple comparison test were used. A p-value < 0.05 was considered statistically significant.

## List of Supplementary Materials

**Table S1. Gene list of significantly expressed in Cm1–10 cardiomyocyte clusters.**

## Acknowledgments

Single nuclei RNA sequencing was done by MedGenome Inc., Foster City, CA.

## Author contributions

Conceptualization: TA, FL

Methodology: TA, RS, RSK, NP, GR, SM, CG, MK, CL, KH,

Investigation: TA, RS, RSK, GR, CL, CG

Funding acquisition: FL, JHS

Project administration: FL, JHS

Supervision: FL, JHS

Writing – original draft: TA, FL

Writing – review & editing: RS, RSK, NP, SM, CG, MK, KH, CL

## Data and materials availability

The snRNA-seq datasets have been deposited in the Gene Expression Omnibus (GEO), accession number GSE292370.

